# dsRNAmax: a Multi-Target Chimeric dsRNA Designer for Safe And Effective Crop Protection

**DOI:** 10.1101/2024.12.15.628581

**Authors:** Stephen J Fletcher, Jai Lawrence, Anne Sawyer, Narelle Manzie, Donald Gardiner, Neena Mitter, Christopher Brosnan

**Affiliations:** Centre for Horticultural Science, Queensland Alliance for Food and Agriculture Innovation, University of Queensland, St Lucia, Queensland, 4072, Australia; Charles Sturt University, New South Wales, Australia

## Abstract

Crop protection is undergoing significant evolution, transitioning toward sustainable approaches that minimise impacts on the environment and human health. Exogenous application of double-stranded RNA (dsRNA) that silences pest or pathogen genes via RNAi (RNA interference) has promise as a safe and effective next generation crop protection platform without the need for genetic modification. However, exogenous dsRNA application at scale presents challenges. Specifically, a single dsRNA sequence needs to balance targeting the standing variation in a target pest or pathogen group against the potential for adverse impacts in a vast array of non-target and beneficial organisms at the application site and broader environment. To address these competing demands, we present dsRNAmax (https://github.com/sfletc/dsRNAmax), a software package that employs *k*-mer based assembly of chimeric dsRNA sequences to target multiple related RNA sequences, to broaden the target spectrum. The package ensures designed dsRNAs have no defined contiguous sequence homology with any off-target sequences, which can range from single transcriptomes through to metagenome sequence data and beyond. The efficacy of this package is demonstrated by a dsRNAmax-designed dsRNA that inhibits multiple root-knot nematode species but not a non-target nematode species, despite its susceptibility to environmental RNAi and high homology of the target gene.

**GRAPHICAL ABSTRACT:** 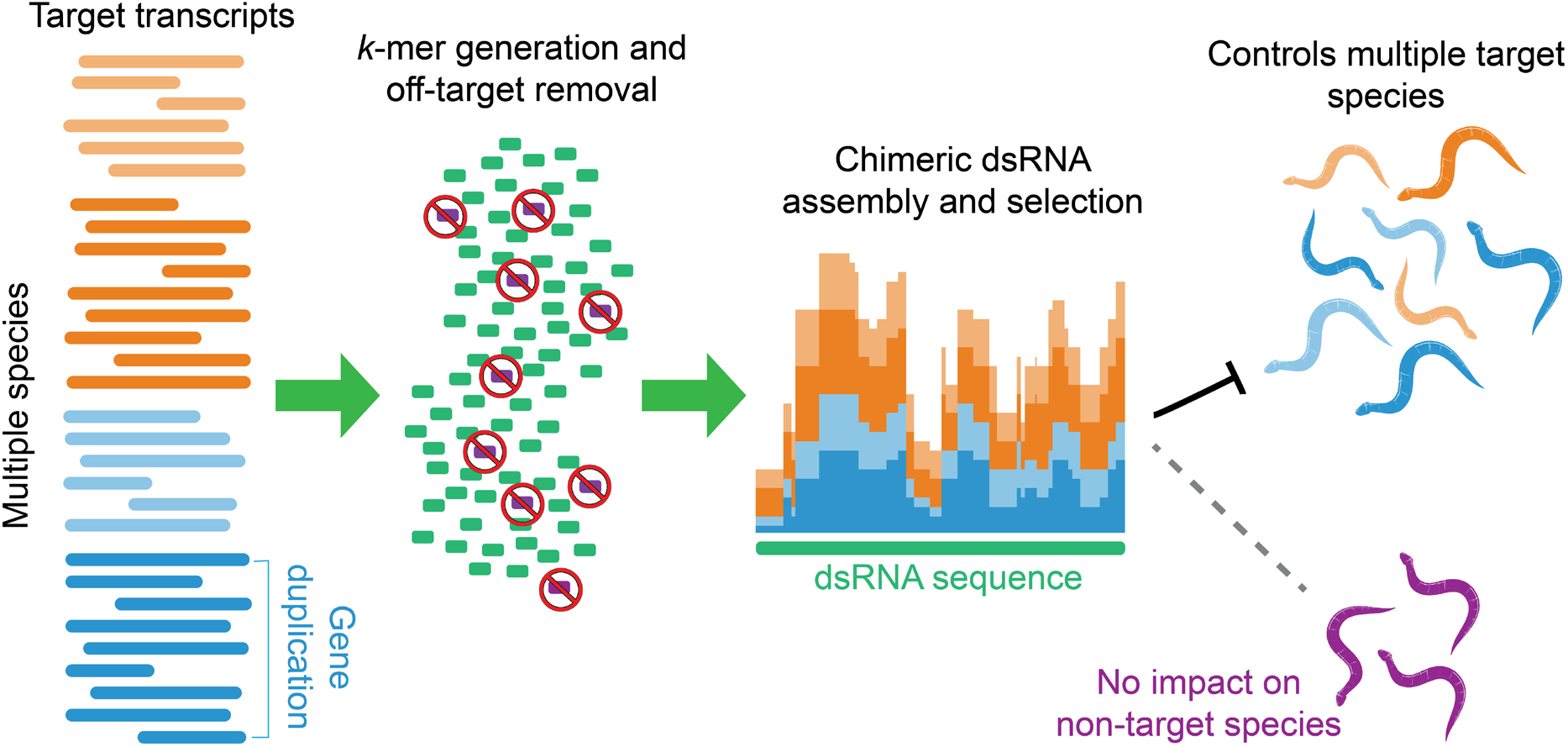

## INTRODUCTION

Pests and pathogens pose a critical threat to global food security, causing annual crop production losses of up to 40% and resulting in worldwide economic damages of approximately US$220 billion (1). Crop protection strategies have historically relied on a combination of host genetic resistance, agronomic management practices, and synthetic pesticides. However, the widespread deployment of synthetic pesticides has raised significant concerns due to their often adverse effects on human health and ecosystem stability, with negative impacts throughout agricultural supply chains (2). In response to these challenges, exogenous RNAi-based crop protection strategies are emerging as promising alternatives. Topical application of dsRNA exploits the cellular mechanism of RNAi to target pest and pathogen species with high specificity, while avoiding many of the regulatory and public acceptance challenges associated with genetic modification (3, 4).

RNAi is a conserved mechanism in most eukaryotes, with roles in virus defence, gene regulation, and maintenance of genome stability. The RNAi pathway is activated by dsRNA, which is processed into 20 to 30 nt small interfering RNAs (siRNAs), depending on the organism (5). When loaded onto ARGONAUTE (AGO) proteins, siRNAs direct cleavage and silencing of complementary RNAs. This pathway can be co-opted via exogenous application of dsRNA to specifically knock down expression of essential pest or pathogen genes, causing mortality in target organisms - a process that is effective against viral, fungal, oomycete, insect and nematode targets (4). A major advantage of this narrow-spectrum approach is that the sequence specificity of RNAi should mitigate unintended impacts on off-target species such as beneficial insects, even if they come into direct contact with the applied dsRNA.

To realise the safety benefits of RNAi-based pesticides, effective design of pest and pathogen targeting dsRNAs is critical. Multiple factors are at play. Firstly, there can be significant genetic variation among target organisms. Variation among viral isolates of the same species serves as a prominent example. Duplication and sequence divergence of functionally comparable target genes can also occur in the same organism. An effective dsRNA must account for this target sequence variation. On the other hand, topical dsRNA application via spraying means potential contact of the dsRNA with a considerable array of non-target organisms, both at the application site and in the surrounding environment. Mitigating unintended impacts, especially when a non-target species is related to the target pest or pathogen, is essential for reaping the environmental and safety benefits RNAi-based biopesticides offer. An example here is avoiding beneficial insects that have been deployed as an IPM strategy whilst targeting specifically a pest. To reduce barriers to adoption including dsRNA synthesis costs and regulatory hurdles relating to multiple active ingredients, application of a single highly efficient dsRNA sequence that targets multiple related pests or pathogens is strongly desired.

Although dsRNA design software packages have been available for decades, they don’t adequately take into consideration the additional requirements that exogenous RNAi -based crop protection bring, including variation amongst sometimes large numbers of target sequences, and complex large-scale off-target requirements. Here, we present dsRNAmax (dsRNA maximizer), an assembly-based chimeric dsRNA designer that maximises the number of dsRNA-derived siRNAs matching target RNAs, while eliminating all contiguous matches to any user-defined off-target sequences, whether they be transcriptomes, genomes or unassembled environmental sample sequences. We used dsRNAmax to design a chimeric dsRNA targeting multiple plant parasitic nematode species but not a designated off-target nematode, and experimentally demonstrated multi-species targeting without off-target impacts.

## MATERIAL AND METHODS

### dsRNAmax development

dsRNAmax is written in Go (Golang), with binaries available for all major operating systems at https://github.com/sfletc/dsRNAmax. It makes use of Go’s concurrency features, efficiently utilising available CPU cores. Memory usage scales with input size, with most dsRNA design tasks able to be completed on commonly available consumer hardware such as laptops and desktops.

### Data input format

dsRNAmax takes inputs for target sequences in FASTA format, and off-target sequences in either FASTA or .kmer format, which is generated using the SeqToKmer package (https://github.com/sfletc/SeqToKmer). When very large off-target datasets are used (for example, paired-end FASTQ files generated from environmental sequencing), the .kmer conversion approach allows for filtering of low count *k*-mers prior to usage, faster execution, and more compact storage. Low count *k*-mers can arise from sequencing errors rather than accurately reflecting underlying sequence variation. When one input sequence is a priority, a bias selection and amount can be specified.

### dsRNAmax framework

For effective dsRNA design based on the ability of processed siRNAs to direct silencing, a *k*-mer approach was adopted (Figure 1). Unique sense-orientation *k*-mers from all input sequences are generated along with a count of input sequences they appear in. The default *k*-mer size of 21 represents the most abundant class of siRNAs in many organisms, although this size is user-selectable. *K*-mers that have a contiguous match to any off-target sequence in either orientation are removed from this pool. One *k*-mer is randomly selected as an initiating seed, then upstream and downstream extension takes place one nucleotide at a time by selecting the most abundant overlapping *k*-mer remaining in the pool, where abundance is measured by the number of input target sequences the *k*-mer exactly matches. Each time a *k*-mer is selected, it is removed from the pool. Where multiple overlapping *k*-mers have an equal abundance, one is randomly selected. Extension in both directions continues until no overlapping *k*-mers are present. If this sequence is longer than the selected dsRNA construct length (default = 300nt), a dsRNA of the defined length is selected by maximising the median of *k*-mer counts matching each target input sequence. Due to inherent random selections, multiple iterations are conducted to identify a more optimal solution (default = 100x).

**Figure 1.**
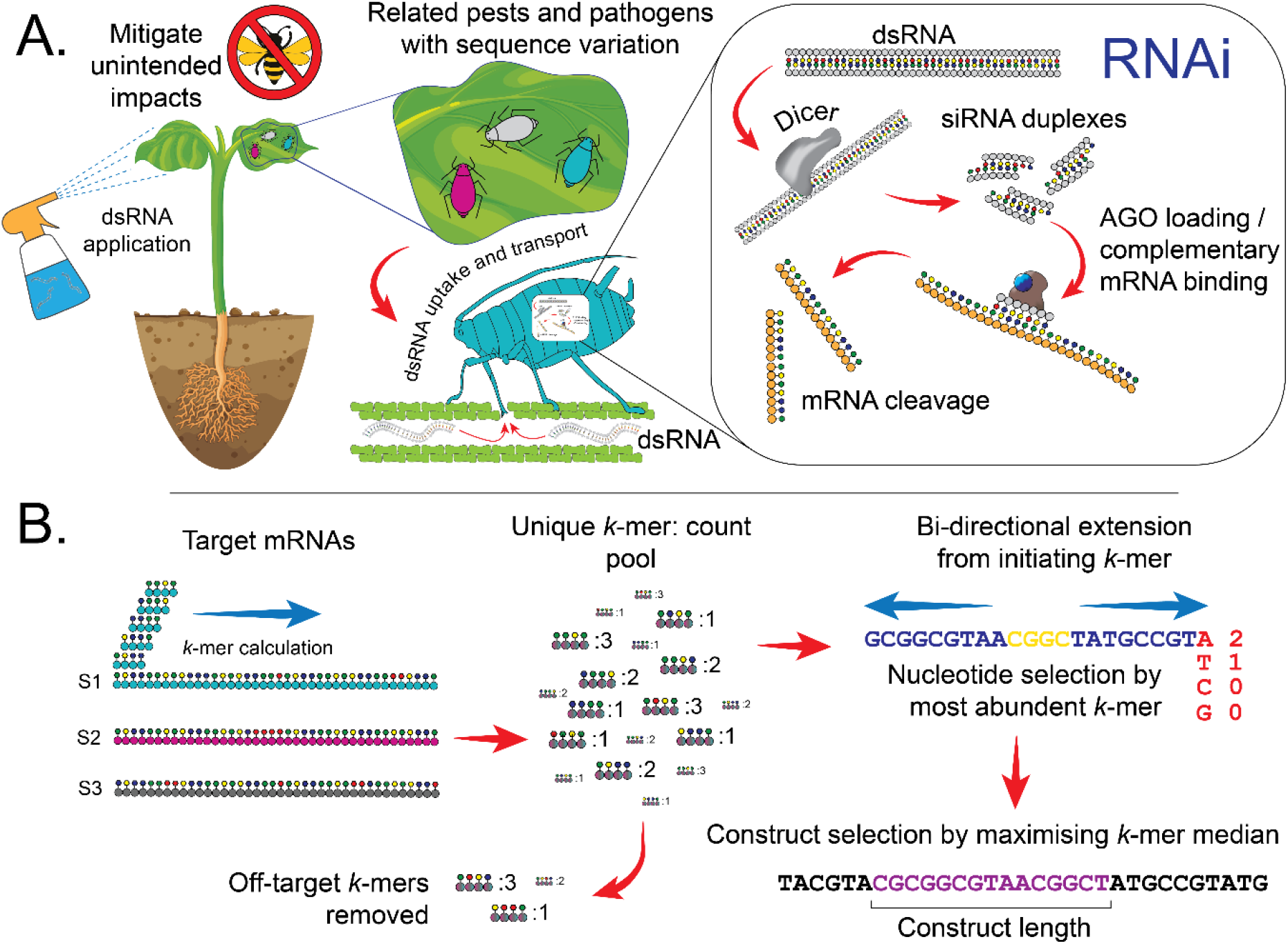
dsRNAmax generates a dsRNA construct targeting multiple sequences whilst mitigating off-target impacts. **A.** Ingestion/uptake of dsRNAs leads to induction of RNAi, where the dsRNA is processed to siRNAs, which are loaded into ARGONAUTES (AGOs) and direct cleavage of complementary host RNAs. **B.** To maximise the number of dsRNA-derived siRNAs that match each target sequence, off-target *k*-mers are first removed, then the remaining *k*-mers assembled by selecting the most abundant for extension, with a dsRNA construct sequence of the desired length selected once assembly is completed.

For each input target sequence, statistics including the number of matching *k*-mers, Smith-Waterman-Gotoh similarity, mean *k*-mer GC content, and matching *k*-mer percentages with 5’U, 5’A and 5’C are presented. dsRNAmax includes the ability to bias the algorithm toward a priority input, which may be desirable where particular biotypes are more important in target population(s). This increases the number of *k*-mers matching the priority sequence but may reduce the median of *k*-mers matching the pool of target sequences. In all cases, no derived *k*-mer in any designed construct will perfectly match any off-target sequence, which is an important consideration for mitigating unintended impacts (6). In situations where this is not possible, no dsRNA will be designed.

### dsRNA design for root-knot nematode (*Meloidogyne*) species

Genome assemblies for four root-knot nematode (RKN) species, *Meloidogyne arenaria, Meloidogyne enterolobii, Meloidogyne incognita* and *Meloidogyne javanica,* were accessed from NCBI (GCA_017562155.1, GCA_903994135.1, GCA_014132215.1 and GCA_900003945.1 respectively) with structural annotations performed using Braker3 (7) and functional annotations via BLAST against UniRef90 (8) and HMMER (www.hmmer.org) against Pfam 37.1(9).

To demonstrate the biological efficacy of a single construct against multiple species in *in vitro* bioassays, we added all *translation elongation factor 1a (TEF1a)* transcript copies annotated in the RKN genomes to a FASTA file as the target input. For the off-target nematode species, *Caenorhabditis elegans* was chosen. The latest genome assembly of this species in FASTA format (GCA_000002985.3) was used as the off-target input. All other dsRNAmax settings except for the off-target setting were left at default. No bias toward any input sequence was used. Two dsRNA sequences were generated, one with the offTarget flag set to 17 (TEF-17) and the other with the offTarget flag set to 21 (TEF-21) (Supplementary Table 1; Supplementary Figure 2).

In addition to generating the dsRNAs for performing on-target and off-target bioassays, we generated two additional dsRNA sense arm sequences to demonstrate the impact of not including any off-target sequences, or using the *C. elegans* off-target genome, but biasing the dsRNA toward one *TEF* sequence, *M. enterolobii* g3046.t1. These example dsRNAs were not used in *in vitro* assays.

### dsRNA generation

Chimeric DNA templates were synthesised by Integrated DNA Technologies. dsRNA was generated using a HiScribe T7 transcription kit (New England Biolabs). For *C. elegans* targeting, dsRNA was synthesised from cDNA generated from N2 mixed stage nematodes. Oligonucleotides used to generate dsRNA are listed in Supplementary Table 2.

#### *Meloidogyne spp.* growth and maintenance

*Meloidogyne spp.* nematodes were provided by Queensland Department of Primary Industries, who maintained colonies on tomato roots (cv Tiny Tim).

#### *Meloidogyne spp.* bioassays

Juvenile 2 (J2) stage *M. incognita, M. javanica* or *M. arenaria* were treated with dsRNA of the indicated sequence and concentration in 50 or 100 µl supplemented with 10 mM octopamine hydrochloride (Sigma Aldrich). Treated nematodes were incubated overnight at room temperature with survival assessed after 24 h under a dissecting microscope.

#### *Caenorhabditis elegans* growth and maintenance

*C. elegans* culture and maintenance were performed using standard techniques(10). Nematodes used were wildtype N2.

#### *Caenorhabditis elegans* bioassays

Assessment of dsRNA targeting *C. elegans* was performed using a brood assay. Synchronised L4 stage worms were incubated in a solution containing 50 ng/µl of the indicated dsRNA (GFP, TEF-17, TEF-21 or ceTEF) for 24 hours. Three nematodes per biological replicate and treatment were transferred to seeded NGM plates and allowed to mature to egg laying adults. These were left for 4 h to lay eggs and then removed. The number of hatched L1/L2 nematodes were then counted.

### Benchmarking dataset

To benchmark dsRNAmax with larger datasets, we downloaded 339 full-length Cucumber Mosaic Virus (CMV) RNA3 genome sequences from NCBI Genbank. This dataset totalled 747,093 nt in length. For a benchmark metagenome off-target dataset, we downloaded paired-end FASTQ files from the NCBI Sequence Read Archive run SRR31016968 (Shotgun metagenome of non-amended cucumber rhizospheres:FORC-inoculated). This run had a total of 7.7G of bases. These reads files were concatenated together, then converted to a .kmer file using SeqToKmer, with the *k*-mer length set to the default of 21 and the minKmerCount filter set to 2. This file was then used as an off-target input to dsRNAmax using the -offTargetKmers flag.

### Statistical analysis

Statistical analysis of RKN kill curve data was performed using Fisher’s Exact tests on 2×2 matrices of alive/deceased counts for each pair-wise comparison. Generated *p*-values were adjusted for multiple comparison using False Discover Ratey (FDR). *C. elegans* brood size tests were performed using pair-wise *t*-tests followed by FDR multiple *p*-value correction (independent broods were derived from three egg-laying adults).

## RESULTS

### Validation study: Targeting four *Meloidogyne* spp. while avoiding *C. elegans*

To demonstrate the ability of dsRNAmax to design single chimeric dsRNA constructs against multiple targets while avoiding unintended impacts on non-target species, we selected four root-knot nematode (RKN) species (*Meloidogyne incognita, Meloidogyne arenaria, Meloidogyne javanica* and *Meloidogyne enterolobii*) as targets. RKNs are small thread-like roundworms ∼0.5 mm in length (11). They invade the roots of many crops including banana, tomato, carrot and other horticultural and ornamental species, causing significant yield loss. *Meloidogyne* species are ideal for use in validation studies due to their availability, likely effectiveness in *in vitro* dsRNA delivery assays, and importance as crop pests (11). *M. enterolobii* was used exclusively for *in silico* design as it is a biosecurity risk in Australia and could not be included in experiments.

We chose *C. elegans* as an off-target species for several reasons. Though not considered a beneficial organism, *C. elegans*’ susceptibility to environmental RNAi and well-defined *in vitro* dsRNA response meant that a dsRNA too closely matching its own transcripts would likely have a negative phenotypic impact, making it an ideal control case for an RKN-targeting chimeric dsRNA.

We reasoned that targeting a constitutively expressed, highly conserved and essential gene would be the best way to test dsRNAmax’s design ability without other potential ectopic effects interfering with our interpretations. For this reason, we chose *Translation Elongation Factor 1a* (*TEF1a*). *TEF1a* is a highly conserved nucleotide-binding protein involved in the elongation of polypeptide chains. *TEF1a* was previously shown to be an effective exogenous dsRNA target in *Austropuccinia psidii* (myrtle rust) (12). This target conservation increases the likelihood of generating a single chimeric dsRNA that targets multiple species but requires careful design to avoid negative impacts on related organisms.

Multiple copies of putative *TEF1a* transcripts were annotated in each RKN species: six copies in *M. arenaria* and *M. incognita* and nine copies in *M. javanica* and *M. enterolobii.* The functional redundancy of these transcripts is unknown, therefore all 30 transcripts were used as inputs to the dsRNAmax design software. As the degree of acceptable homology to *C. elegans* transcripts was also unknown, we designed two dsRNA constructs, selecting either 21 nt (TEF-21) or 17 nt (TEF-17) as the cut-offs for the maximum contiguous homology to the *C. elegans* genome. As expected, by reducing the allowable contiguous homology to the *C. elegans* genome from 21 nt to 17 nt, the median number of 21 nt matches to all *TEF1a Meloidogyne spp.* transcripts was also reduced (from 200.5 to 185) (Figure 2 A and B). Pairwise alignments of each construct to the *C. elegans EF1-a* transcript are shown in Supplementary Figure 1, with aligned region homologies of 79.3% and 85.1% respectively.

**Figure 2.**
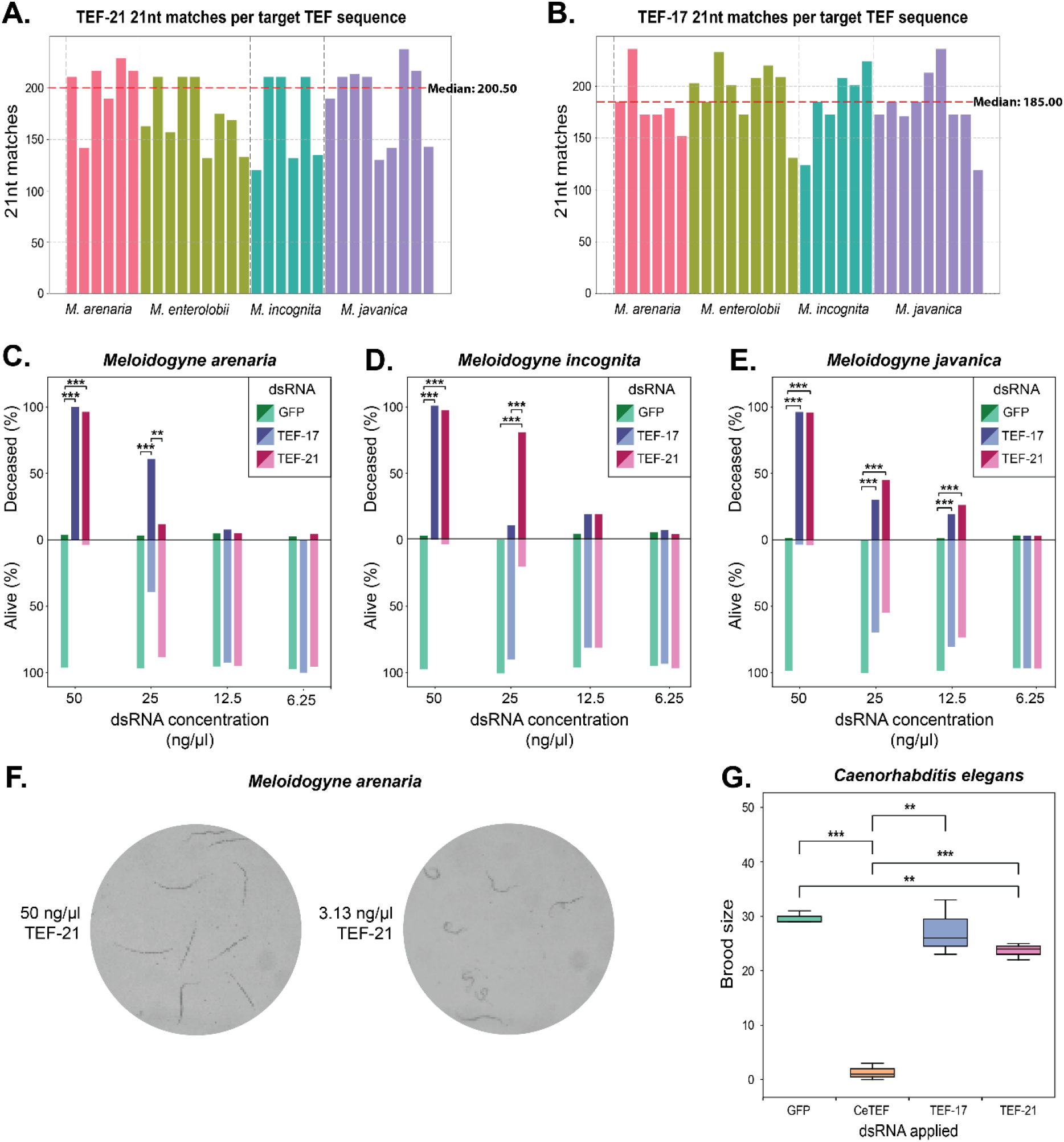
A single 300 nt dsRNA targeting *TEF1a* can control three root-knot nematode species without impacting the nematode *C. elegans*. **A.** Number of 21 nt matches to each *TEF1a* transcript when the maximum contiguous match to the defined off-target *C. elegans* genome is 20 nt (median 21 nt matches of the dsRNA to all targets = 200.5). **B.** Number of 21 nt matches to each *TEF1a* transcript when the maximum contiguous match to the defined off-target *C. elegans* genome is 16 nt (median 21 nt matches of the dsRNA to all targets = 185). **C.** dsRNA concentration-kill curve for *M. arenaria* **D.** dsRNA concentration-kill curve for *M. incognita* **E.** dsRNA concentration-kill curve for *M. javanica*. **F.** Example images of deceased (left) and alive *M. arenaria* (right) nematodes treated with 50 ng/µl and 3.18 ng/µl of TEF-21 dsRNA respectively. **G.** *C. elegans* brood size in response to dsRNA application (50 ng/µl). Kill curve statistical tests were performed using Fisher’s Exact Test followed by FDR multiple *p*-value correction. Brood size statistic tests were performed using pair-wise *t*-tests followed by FDR multiple *p*-value correction (independent broods were derived from three egg-laying adults).

To test the ability of our software to specifically target multiple RKN species, we treated three selected species, *M. arenaria, M. incognita* and *M. javanica,* with varying concentrations of TEF-21 and TEF-17 dsRNAs, along with a non-target/negative Green Fluorescent Protein (GFP) dsRNA control, in *in vitro* soaking assays. After 24 hours of soaking, RKNs were scored for alive and deceased counts on plates. Kill curves for each RKN species using a two-fold dilution series of dsRNA concentrations from 50 ng/µl to 6.25 ng/µl are shown in Figure 2 C-E. Example images of alive and deceased morphology are shown in Figure 2 F. A concentration of 50 ng/µl of either RKN targeting dsRNA was required for complete mortality in all RKN species (Figure 2 C-E, Supplementary Tables 3-5). Concentrations of 25 ng/µl and 12.5 ng/µl had partial levels of mortality, while concentrations of 6.25 ng/µl were ineffective. The *GFP*-targeting dsRNA had no negative impact on RKN health, demonstrating that TEF-21 and TEF-17 were acting in an RNAi-dependent sequence specific manner.

An important design factor in this example is the avoidance of negative impacts on our selected off-target species, *C. elegans*. For the off-target assays, a positive control dsRNA exactly matching the *C. elegans EF1-a* transcript was used (CeTEF). None of the dsRNAs had negative impacts on *C. elegans* mortality upon soaking, but the impact on the subsequent generation as evidenced by brood size was clear (Figure 2 G). The CeTEF dsRNA resulted in near zero brood sizes when compared with the negative control GFP dsRNA treated samples and TEF-17 and TEF-21 treatments. Interestingly, there was a significant difference between the *GFP* non-target control and the TEF-21 treatment, with a very slight decrease in brood size. This was not seen with the TEF-17 treated samples, matching the stringency of the targeting predictions. While there was no significant difference between TEF-21 and TEF-17 dsRNA impacts on *C. elegans* brood size, the individual comparisons to the negative GFP control trend with our *in silco* stringency predictions.

### Impact of off-target and bias selections

In general, adding off-target sequences reduces the median number of *k*-mers matching each target input sequence if off-target *k*-mers are derived from related species and/or target genes are highly conserved. By reducing the pool of *k*-mers from which the pre-dsRNA sequences can be assembled, the likelihood increases that less abundant *k*-mers must be assembled. For example, if the *k*-mer that matches the most target input sequences is removed, the second most abundant overlapping *k*-mer is used for extension. Where the removal of a *k*-mer means that extension can no longer take place, a different region of the assembled sequence is used for the final chimeric dsRNA selection. Design against the *Meloidogyne spp. TEF1a* transcript set without the use of off-targets generated a median number of 21 nt matches to each transcript of 203 out of a maximum of 280 sense orientation *k*-mers for a 300 nt construct (Supplementary Figure 3). Addition of the *C. elegans* genome as an off-target dataset, and an off-target cutoff of 17 (i.e. there are no matches ≥17 nt between the dsRNA and the off-target dataset in either orientation), resulted in the median 21 nt matches to each target dropping to 185, with 1,305 *k*-mers removed from the initial 8,902 target pool (Supplementary Figure 2).

A further consideration of multi-target dsRNA design is the relative importance of individual targets within the target dataset. In some situations, a certain viral isolate or pathogen species may be considered of greatest importance for control. Given that a range of *k*-mer matches between the designed dsRNA construct and the target dataset may exist, biasing the design can ensure that a specified target sequence sits toward the top of this distribution. Biasing in this way can however reduce the median dsRNA-derived *k*-mer matches to the overall target dataset.

For chimeric dsRNA design against 30 *Meloidogyne spp. TEF1a* transcripts with a 17 nt *C. elegans* off-target selection, dsRNA derived 21 nt matches to these targets ranged from 119 to 236, with the median 185 (Supplementary Figure 2). The number of 21 nt matches to the specific *M. enterolobii* transcript g3046.t1 in this situation was 203. By biasing toward this transcript by flagging it and setting the biasLvL flag to 10 (which adds 10 copies of this transcript to the design phase), the number of matching dsRNA-derived 21 nt *k*-mers rose to 236, however the median *k*-mer hits reduced to 162 (Supplementary Figure 4). That is, the increase in 21 nt matches to g3046.t1 comes at the cost of reduced matches to all transcripts.

### dsRNAmax large dataset performance

Benchmarking on larger input and off-target datasets showed that dsRNAmax can rapidly design chimeric dsRNAs using desktop hardware (Supplementary Table 6). A larger target input set including 339 Cucumber Mosaic Virus (CMV) RNA segments was used, along with an off-target .kmer dataset generated from cucumber soil rhizosphere metagenome sequencing. All design runs took less than 20 seconds. As FASTQ files can’t directly be used, the conversion to .kmer format provides a highly performant alternative in which off-target *k*-mers are stored as unsinged 64-bit integers. In this instance, singleton *k*-mers were removed to reduce the likelihood of sequencing errors impacting the off-target *k*-mer removal process. Similar to index generation for read alignment software, this process can be slow (over 10 minutes for a 7.7 GB read dataset), but only needs to be carried out once for each off-target dataset and is largely independent of on-target design concerns. The only requirement for the off-target contiguous match length is that it be less than or equal to the on-target *k*-mer size.

## DISCUSSION

Through validation experiments on major crop pests, root-knot nematodes, we have shown that the dsRNAmax software successfully achieves its primary objective: designing effective chimeric dsRNA constructs targeting multiple pest species while avoiding significant impacts on non-target organisms. By leveraging the conserved nature of the *TEF1a* gene across the four *Meloidogyne* species, despite multiple copies being present in each genome, dsRNAmax generated a single chimeric dsRNA construct that elicited high mortality in three tested species, confirming its control efficacy.

The use of the off-target *C. elegans* genome demonstrated the software’s capacity to minimise unintended impacts. Constructs designed with stricter homology thresholds (e.g., TEF-17) demonstrated no significant difference in brood size to the non-target GFP control. In contrast, there was a significant but small magnitude difference between TEF-21 and GFP. If this were a real-world example and the off-target species were beneficial predatory nematodes present at application, the more conservative 17 nt off-target selection would be chosen to ensure no negative impacts on these beneficial species.

This work highlights the flexibility of dsRNAmax in balancing specificity and efficacy through adjustable parameters, such as homology thresholds and target prioritisation. The observed trade-off between maximizing coverage across all target transcripts and biasing the design toward specific sequences underscores the software’s utility for tailoring dsRNA constructs to specific pest management needs.

Existing dsRNA design packages tend to focus on optimising a selected region of a single RNA for targeting, based on factors such as AGO loading preference, siRNA strand selection and accessibility of target sites (Supplementary Table 7). dsRNAmax is designed for use against a wide range of targets, including plant viruses, fungi, oomycetes, nematodes and insects, each with potentially differing siRNA biogenesis and AGO loading strategies. It is also intended for application in variable field conditions where other biotic and abiotic stresses may be present. Relatively little is known about dsRNA processing outside of model organisms in controlled laboratory conditions. Data concerning exogenous dsRNA uptake and processing is even more limited. To address these challenges, the primary design rule is to maximise the number of siRNAs that perfectly match each input sequence, rather than attempt to fit the preferences of a target species. Given that in some instances these preferences may be known (13–15), the software output includes information such as GC content and 5’ ribonucleotide occurrence for the matching *k*-mers for each target input, which may help guide design parameters.

### Multi-target design and sequence conservation challenges

The development of dsRNAmax software for pest and pathogen control presents several critical considerations. Multi-target chimeric dsRNA design can be effective in scenarios involving conserved genetic targets, but limitations exist when functionally similar targets become too divergent at the transcript sequence level. This necessitates a nuanced approach to dsRNA design that recognises the complexity of genetic variation. A relevant example of this divergence is plant virus genomes of the same species, where significant sequence diversity can occur in some instances. Accordingly, generating effective universal chimeric dsRNAs as target sequences diverge becomes increasingly difficult. In such situations, multiple or concatenated dsRNAs may be the best solution.

### Risk assessment and practical implications

While off-target effects represent a complex consideration in dsRNA design, our experimental nematode data indicate that multiple species can be targeted with a single dsRNA and negative impacts in other dsRNA-susceptible species can be avoided. Bioinformatics-based design however should be understood as a preliminary step rather than a definitive solution. Rigorous empirical testing remains essential, particularly when targeting novel or less-studied species, both for on-target efficacy and off-target avoidance. The cost and time requirements of synthesising chimeric dsRNAs should also be considered when screening potential target genes. In some instances, pre-screening targets with perfectly matching dsRNAs may be desirable.

## Conclusion

Effective dsRNA design requires a multi-layered approach that balances computational prediction with empirical validation. The most successful strategies will integrate bioinformatics with experimental testing, particularly when addressing genetically varied targets in less-studied species and complex ecological systems. dsRNAmax offers a software solution that sits in the design phase of this workflow, generating safe and specific universal chimeric dsRNAs for use in crop pest and pathogen control applications.

## DATA AVAILABILITY

All data are incorporated into the article and its online supplementary material.

## SUPPLEMENTARY DATA

Supplementary Data are available online

## AUTHOR CONTRIBUTIONS

Stephen Fletcher: Conceptualisation, Software, Formal analysis, Methodology, Validation, Writing-original draft. Jai Lawrence: Investigation, Validation. Anne Sawyer: Investigation, Methodology, Writing – review & editing. Narelle Manzie: Project administration, Writing – review & editing. Donald Gardiner: Project administration, Writing – review & editing, Neena Mitter: Funding acquisition, Project administration, Writing – review & editing. Christopher Brosnan: Conceptualisation, Methodology, Validation, Writing-original draft.

## ACKNOWLEDGEMENTS

We would like to thank Jennifer Coban, Wayne O’Neill, Tim Shuey and Dylan Corner from Queensland Department of Primary Industries for their technical advice and for providing the root knot nematodes.

## FUNDING

This work was supported by Australian Research Council [IH190100022]. Funding for open access charge: Australian Research Council. AS was supported by an Advance Queensland Industry Research Fellowship.

## CONFLICT OF INTEREST

The authors report no conflict of interest.

## Notes

### Competing Interest Statement

The authors have declared no competing interest.

