## Supplementary Data for "dsRNAmax: a Multi-Target Chimeric dsRNA Designer for Safe And Effective Crop Protection"

**Supplementary Table 1: *Meloidogyne* universal dsRNAs with *C. elegans* off-target avoidance.**

| dsRNA name | dsRNA sense arm sequence (5′ to 3′) |
|---|---|
| ***TEF-17*** | TCAAGAACATGATTACTGGTACATCCCAAGCTGATTGTGCTGTTTTGGTTGTTG CTTGTGGTACTGGAGAGTTCGAGGCTGGAATTTCCAAGAACGGTCAAACTCGC GAGCATGCTCTTCTCGCTCAGACTTTGGGAGTGAAGCAACTTATCGTTGCCTGT AATAAGAAGATTGGTTACAACCCTGCAACCGTTGCTTTCGTTCCTATTTCTGGCT TTAATGGCGACAATATGTTGGAGCCGTCTGATAAGATGCCTTGGTTCAAAGGA TGGGCTATTGAAAGGAAGGATGGAAATGCTA |
| ***TEF-21*** | AAATACTATGTCACAATTATCGATGCTCCTGGACATCGTGACTTCATCAAGAAC ATGATTACTGGTACATCCCAAGCTGATTGTGCTGTTTTGGTTGTTGCTTGTGGTA CTGGAGAGTTCGAGGCTGGAATTTCTAAGAATGGTCAAACTCGCGAGCATGCT CTTCTCGCTCAGACTTTGGGAGTGAAGCAACTTATTGTTGCGTGTAATAAGATG GACACGACTGAGCCACCATTTTCGGAAGCTCGTTTTGAGGAAGTTAAGAATGA AGTCTCCAGCTTTATTAAGAAGATTGGTTAC |

**Supplementary Table 2: PCR primers used for amplifying dsRNA templates**

| Primer name | Sequence (5’ to 3’) |
|---|---|
| **CB-c.el-TEF-T7-F1** | TAATACGACTCACTATAGGGAAGTACTACATCACCATCATCGATGC |
| **CB-c.el-TEF-T7-R1** | TAATACGACTCACTATAGGGGTATCCGATCTTCTTGATGAATCCAG |
| **CB-mu-21nt-TEF-F** | TAATACGACTCACTATAGGGAAATACTATGTCACAATTATCG |
| **CB-mu-21nt-TEF-R** | TAATACGACTCACTATAGGGGTAACCAATCTTCTTAATAAAGC |
| **CB-mu-17nt-TEF-F** | TAATACGACTCACTATAGGGTCAAGAACATGATTACTGGTAC |
| **CB-mu-17nt-TEF-R** | TAATACGACTCACTATAGGGTAGCATTTCCATCCTTCCTTTC |

**Supplementary Table 3: Alive and deceased raw count data for *M. arenaria*.**

| dsRNA (ng/ul) | GFP_alive | GFP_dead | Tef_17_alive | Tef_17_dead | Tef_21_alive | Tef_21_dead |
|---|---|---|---|---|---|---|
| **50** | 26 | 1 | 0 | 22 | 1 | 26 |
| **25** | 29 | 1 | 9 | 14 | 15 | 2 |
| **12.5** | 40 | 2 | 24 | 2 | 19 | 1 |
| **6.25** | 37 | 1 | 25 | 0 | 21 | 1 |
| **3.125** | 19 | 1 | 15 | 0 | 30 | 1 |
| **1.5625** | 34 | 1 | 25 | 0 | 12 | 1 |

**Supplementary Table 4: Alive and deceased raw count data for *M. incognita*.**

| dsRNA (ng/ul) | GFP_alive | GFP_dead | Tef_17_alive | Tef_17_dead | Tef_21_alive | Tef_21_dead |
|---|---|---|---|---|---|---|
| **50** | 32 | 1 | 0 | 18 | 1 | 28 |
| **25** | 13 | 0 | 17 | 2 | 6 | 24 |
| **12.5** | 22 | 1 | 17 | 4 | 17 | 4 |
| **6.25** | 17 | 1 | 26 | 2 | 24 | 1 |
| **3.125** | 11 | 0 | 21 | 0 | 13 | 0 |
| **1.5625** | 27 | 1 | 11 | 1 | 14 | 1 |

**Supplementary Table 5: Alive and deceased raw count data for *M. javanica*.**

| dsRNA (ng/ul) | GFP_alive | GFP_dead | Tef_17_alive | Tef_17_dead | Tef_21_alive | Tef_21_dead |
|---|---|---|---|---|---|---|
| **50** | 67 | 1 | 2 | 58 | 3 | 75 |
| **25** | 61 | 0 | 50 | 22 | 29 | 24 |
| **12.5** | 68 | 1 | 53 | 13 | 47 | 17 |
| **6.25** | 53 | 2 | 57 | 2 | 60 | 2 |
| **3.125** | 60 | 1 | 84 | 2 | 60 | 1 |
| **1.5625** | 61 | 1 | 52 | 1 | 0 | 0 |

**TEF-21**

```
TEF-21     AAATACTATGTCACAATTATCGATGCTCCTGGACATCGTGACTTCATCAAGAACATGATT  60
           || |||||  |||| ||||| ||||| || |||||||||||| |||||||||||||||||
Ce-EF1a    AAGTACTACATCACCATTATTGATGCCCCAGGACATCGTGATTTCATCAAGAACATGATT  332

TEF-21     ACTGGTACATCCCAAGCTGATTGTGCTGTTTTGGTTGTTGCTTGTGGTACTGGAGAGTTC  120
           || || || |||||||| ||||| || |||||||| || || || || || |||||||||
Ce-EF1a    ACCGGAACCTCCCAAGCCGATTGCGCCGTTTTGGTCGTCGCCTGCGGAACCGGAGAGTTC  392

TEF-21     GAGGCTGGAATTTCTAAGAATGGTCAAACTCGCGAGCATGCTCTTCTCGCTCAGACTTTG  180
           ||||| ||||| || ||||| || ||||| |||||||||||||| || || |||||  | 
Ce-EF1a    GAGGCCGGAATCTCCAAGAACGGACAAACCCGCGAGCATGCTCTCCTTGCCCAGACCCTT  452

TEF-21     GGAGTGAAGCAACTTATTGTTGCGTGTAATAAGATGGACACGACTGAGCCACCATTTTCG  240
           ||||| ||||| || || || || || || ||||||||| | || ||||||||||| || 
Ce-EF1a    GGAGTCAAGCAGCTCATCGTCGCTTGCAACAAGATGGACTCCACCGAGCCACCATTCTCT  512

TEF-21     GAAGCTCGTTTTGAGGAAGTTAAGAATGAAGTCTCCAGCTTTATTAAGAAGATTGGTTAC  300
           || ||||||||    ||  | |  || || |||||| | || || |||||||| || |||
Ce-EF1a    GAGGCTCGTTTCACTGAGATCACCAACGAGGTCTCCGGATTCATCAAGAAGATCGGATAC  572
```

**TEF-17**

```
TEF-17     TCAAGAACATGATTACTGGTACATCCCAAGCTGATTGTGCTGTTTTGGTTGTTGCTTGTG  60
           ||||||||||||| ||||| |||||||| || ||||| || || ||||| || ||||| |
Ce-EF1a    TCAAGAACATGATCACTGGAACATCCCAGGCCGATTGCGCCGTCTTGGTCGTCGCTTGCG  387

TEF-17     GTACTGGAGAGTTCGAGGCTGGAATTTCCAAGAACGGTCAAACTCGCGAGCATGCTCTTC  120
           | |||||||| |||||||| ||||| ||||||||||| |||||||||||||| |||||||
Ce-EF1a    GAACTGGAGAATTCGAGGCCGGAATCTCCAAGAACGGACAAACTCGCGAGCACGCTCTTC  447

TEF-17     TCGCTCAGACTTTGGGAGTGAAGCAACTTATCGTTGCCTGTAATAAGA  168
           | || |||||  | ||||| |||||||| ||||||||||| || ||||
Ce-EF1a    TTGCCCAGACCCTTGGAGTCAAGCAACTCATCGTTGCCTGCAACAAGA  495
```

**Supplementary Figure 1: Pairwise alignments between the TEF-21 (top) and TEF-17 (bottom) dsRNA sense arms to *C. elegans EF1a*.** Mismatches are shaded in orange. The percentage homology of the aligned regions is 79.3% and 85.1% respectively.

```
2024/12/10 13:50:31 dsRNAmax - dsRNA maximizer (Version: 1.1.12)
2024/12/10 13:50:31 Target FASTA File: ../ref/TEF.fa
2024/12/10 13:50:31 Off-target FASTA File: ../ref/GCF_000002985.6_WBcel235_genomic.fasta
2024/12/10 13:50:31 Loading target sequences...
     ---> 30 sequences loaded
2024/12/10 13:50:31 Getting target sequence kmers...
2024/12/10 13:50:31 8,902 target kmers loaded
2024/12/10 13:50:31 Removing off-target kmers from FASTA files...
Total off-target-matching kmers removed: 1305

2024/12/10 13:50:32 Finding best construct...

Results:
+-------------------------------------------------------------+-------------+--------------------+------------------+---------+---------+---------+
|                    TARGET SEQUENCE HEADER                   | 21NT MATCHES | SWG SIMILARITY (%) | KMER MEAN GC (%) | 5'U (%) | 5'A (%) | 5'C (%) |
+-------------------------------------------------------------+-------------+--------------------+------------------+---------+---------+---------+
| g3046.t1__seq_id=CAJEWN010000016.1_type=cds_species=Menterolobii  | 203 | 53.3 | 46.2 | 24.1 | 30.5 | 28.6 |
| g17458.t1__seq_id=CAJEWN010000248.1_type=cds_species=Menterolobii | 209 | 53.3 | 46.3 | 23.4 | 30.6 | 28.7 |
| g4219.t1__seq_id=BLLR01000009.1_type=cds_species=Mincognita       | 224 | 94.7 | 45.5 | 24.1 | 30.4 | 28.1 |
| g8398.t1__seq_id=CEWN01001434.1_type=cds_species=Mjavanica        | 173 | 52.0 | 45.7 | 24.3 | 30.1 | 28.9 |
| g16315.t1__seq_id=CEWN01003635.1_type=cds_species=Mjavanica       | 171 | 53.0 | 47.1 | 22.8 | 31.0 | 29.2 |
| g2430.t1__seq_id=JAEEAS010000070.1_type=cds_species=Marenaria     | 173 | 52.0 | 45.7 | 24.3 | 30.1 | 28.9 |
| g7704.t1__seq_id=CEWN01001282.1_type=cds_species=Mjavanica        | 213 | 52.3 | 46.7 | 20.2 | 33.3 | 28.6 |
| g26932.t1__seq_id=CEWN01007834.1_type=cds_species=Mjavanica       | 173 | 52.0 | 45.7 | 24.3 | 30.1 | 28.9 |
| g2400.t1__seq_id=JAEEAS010000070.1_type=cds_species=Marenaria     | 179 | 52.0 | 46.8 | 21.2 | 33.0 | 26.8 |
| g36955.t1__seq_id=CAJEWN010002398.1_type=cds_species=Menterolobii | 220 | 56.0 | 47.1 | 24.1 | 29.1 | 27.7 |
| g28756.t1__seq_id=CAJEWN010000898.1_type=cds_species=Menterolobii | 131 | 51.0 | 45.5 | 28.2 | 29.0 | 27.5 |
| g47475.t1__seq_id=JAEEAS010001409.1_type=cds_species=Marenaria    | 152 | 51.0 | 46.1 | 27.0 | 27.6 | 27.6 |
| g11711.t1__seq_id=CEWN01002262.1_type=cds_species=Mjavanica       | 173 | 52.0 | 47.1 | 24.9 | 28.9 | 27.7 |
| g9315.t1__seq_id=CAJEWN010000078.1_type=cds_species=Menterolobii  | 173 | 52.0 | 47.1 | 24.9 | 28.9 | 27.7 |
| g17491.t1__seq_id=CAJEWN010000248.1_type=cds_species=Menterolobii | 208 | 53.3 | 47.5 | 22.1 | 29.8 | 28.8 |
| g1759.t1__seq_id=JAEEAS010000058.1_type=cds_species=Marenaria     | 173 | 52.0 | 47.1 | 24.9 | 28.9 | 27.7 |
| g11578.t1__seq_id=BLLR01000031.1_type=cds_species=Mincognita      | 173 | 52.0 | 47.1 | 24.9 | 28.9 | 27.7 |
| g4252.t1__seq_id=BLLR01000009.1_type=cds_species=Mincognita       | 208 | 53.3 | 47.5 | 22.1 | 29.8 | 28.8 |
| g42043.t1__seq_id=CEWN01017646.1_type=cds_species=Mjavanica       | 236 | 54.0 | 46.7 | 22.0 | 30.9 | 28.0 |
| g9360.t1__seq_id=CAJEWN010000078.1_type=cds_species=Menterolobii  | 201 | 53.0 | 46.2 | 23.9 | 30.3 | 28.4 |
| g27345.t1__seq_id=JAEEAS010000752.1_type=cds_species=Marenaria    | 236 | 54.0 | 46.7 | 22.0 | 30.9 | 28.0 |
| g11539.t1__seq_id=BLLR01000031.1_type=cds_species=Mincognita      | 201 | 53.0 | 46.2 | 23.9 | 30.3 | 28.4 |
| g4100.t1__seq_id=CEWN01000556.1_type=cds_species=Mjavanica        | 185 | 53.0 | 46.2 | 22.2 | 31.9 | 27.0 |
| g34890.t1__seq_id=CAJEWN010001880.1_type=cds_species=Menterolobii | 233 | 55.0 | 46.9 | 22.3 | 30.5 | 27.5 |
| g1731.t1__seq_id=JAEEAS010000058.1_type=cds_species=Marenaria     | 185 | 53.0 | 46.2 | 22.2 | 31.9 | 27.0 |
| g25165.t1__seq_id=CEWN01007035.1_type=cds_species=Mjavanica       | 185 | 53.0 | 46.2 | 22.2 | 31.9 | 27.0 |
| g3081.t1__seq_id=CAJEWN010000016.1_type=cds_species=Menterolobii  | 185 | 53.0 | 46.2 | 22.2 | 31.9 | 27.0 |
| g18725.t1__seq_id=BLLR01000069.1_type=cds_species=Mincognita      | 185 | 53.0 | 46.2 | 22.2 | 31.9 | 27.0 |
| g18693.t1__seq_id=BLLR01000069.1_type=cds_species=Mincognita      | 124 | 50.0 | 43.9 | 25.0 | 29.8 | 29.0 |
| g8399.t1__seq_id=CEWN01001434.1_type=cds_species=Mjavanica        | 119 | 45.7 | 44.5 | 26.1 | 27.7 | 28.6 |
+-------------------------------------------------------------+-------------+--------------------+------------------+---------+---------+---------+

Median of kmer hits to each target sequence: 185

dsRNA sense-arm sequence - 45.0% GC content
TCAAGAACATGATTACTGGTACATCCCAAGCTGATTGTGCTGTTTTGGTTGTTGCTTGTGGTACTGGAGAGTTCGAGGCTGGAATTTCCAAGAACGGTCAAACTCGCGAGCATGCTCTTCTCGCTCAGACTTTGGGAGTGAAGCAACTTATCGTTGCCTGTAATA
AGAAGATTGGTTACAACCCTGCAACCGTTGCTTTCGTTCCTATTTCTGGCTTTAATGGCGACAATATGTTGGAGCCGTCTGATAAGATGCCTTGGTTCAAAGGATGGGCTATTGAAAGGAAGGATGGAAATGCTA
```

**Supplementary Figure 2: Universal *Meloidogyne* dsRNA with the *C. elegans* genome off-target dataset (TEF-17).** 30 *Meloidogyne spp. TEF1a* transcript sequences were used as target inputs. The *C. elegans* genome assembly (WBcel235) served as the off-target input in FASTA format, with the off-target cutoff set to 17. All other options were at the default settings (on-target *k*-mer length = 21, construct length = 300, 100 iterations). The final design had a median of 185 21 nt *k*-mer matchers to each target TEF sequence.

```
2024/12/10 13:53:20 dsRNAmax - dsRNA maximizer (Version: 1.1.12)
2024/12/10 13:53:20 Target FASTA File: ../ref/TEF.fa
2024/12/10 13:53:20 Loading target sequences...
    ---> 30 sequences loaded
2024/12/10 13:53:20 Getting target sequence kmers...
2024/12/10 13:53:20 8,902 target kmers loaded
2024/12/10 13:53:20 Finding best construct...

Results:
+--------------------------------------------------------------+-------------+--------------------+------------------+---------+---------+---------+
|                      TARGET SEQUENCE HEADER                  | 21NT MATCHES | SWG SIMILARITY (%) | KMER MEAN GC (%) | 5'U (%) | 5'A (%) | 5'C (%) |
+--------------------------------------------------------------+-------------+--------------------+------------------+---------+---------+---------+
| g3046.t1__seq_id=CAJEWN010000016.1_type=cds_species=Menterolobii  | 224 | 95.0 | 47.0 | 24.6 | 29.9 | 25.9 |
| g17458.t1__seq_id=CAJEWN010000248.1_type=cds_species=Menterolobii | 224 | 95.0 | 47.0 | 24.6 | 29.9 | 25.9 |
| g4219.t1__seq_id=BLLR01000009.1_type=cds_species=Mincognita       | 224 | 95.0 | 47.0 | 24.6 | 29.9 | 25.9 |
| g8398.t1__seq_id=CEWN01001434.1_type=cds_species=Mjavanica        | 206 | 95.0 | 45.3 | 24.3 | 31.1 | 27.2 |
| g16315.t1__seq_id=CEWN01003635.1_type=cds_species=Mjavanica       | 218 | 95.0 | 46.8 | 22.5 | 30.7 | 28.0 |
| g2430.t1__seq_id=JAEEAS010000070.1_type=cds_species=Marenaria     | 206 | 95.0 | 45.3 | 24.3 | 31.1 | 27.2 |
| g7704.t1__seq_id=CEWN01001282.1_type=cds_species=Mjavanica        | 213 | 95.3 | 47.6 | 23.5 | 30.0 | 25.8 |
| g26932.t1__seq_id=CEWN01007834.1_type=cds_species=Mjavanica       | 203 | 94.0 | 45.5 | 24.1 | 31.0 | 26.6 |
| g2400.t1__seq_id=JAEEAS010000070.1_type=cds_species=Marenaria     | 238 | 98.0 | 46.1 | 23.9 | 30.7 | 26.9 |
| g36955.t1__seq_id=CAJEWN010002398.1_type=cds_species=Menterolobii | 221 | 95.0 | 48.0 | 24.4 | 29.9 | 25.8 |
| g28756.t1__seq_id=CAJEWN010000898.1_type=cds_species=Menterolobii | 164 | 91.0 | 45.6 | 26.2 | 30.5 | 26.2 |
| g47475.t1__seq_id=JAEEAS010001409.1_type=cds_species=Marenaria    | 182 | 93.0 | 45.9 | 26.4 | 29.1 | 25.3 |
| g11711.t1__seq_id=CEWN01002262.1_type=cds_species=Mjavanica       | 187 | 94.0 | 47.1 | 24.1 | 32.1 | 26.2 |
| g9315.t1__seq_id=CAJEWN010000078.1_type=cds_species=Menterolobii  | 187 | 94.0 | 47.1 | 24.1 | 32.1 | 26.2 |
| g17491.t1__seq_id=CAJEWN010000248.1_type=cds_species=Menterolobii | 186 | 93.3 | 48.0 | 22.6 | 30.1 | 26.3 |
| g1759.t1__seq_id=JAEEAS010000058.1_type=cds_species=Marenaria     | 187 | 94.0 | 47.1 | 24.1 | 32.1 | 26.2 |
| g11578.t1__seq_id=BLLR01000031.1_type=cds_species=Mincognita      | 187 | 94.0 | 47.1 | 24.1 | 32.1 | 26.2 |
| g4252.t1__seq_id=BLLR01000009.1_type=cds_species=Mincognita       | 186 | 93.3 | 48.0 | 22.6 | 30.1 | 26.3 |
| g42043.t1__seq_id=CEWN01017646.1_type=cds_species=Mjavanica       | 207 | 94.3 | 48.0 | 23.7 | 29.5 | 26.6 |
| g9360.t1__seq_id=CAJEWN010000078.1_type=cds_species=Menterolobii  | 239 | 96.3 | 46.1 | 24.7 | 29.7 | 26.4 |
| g27345.t1__seq_id=JAEEAS010000752.1_type=cds_species=Marenaria    | 207 | 94.3 | 48.0 | 23.7 | 29.5 | 26.6 |
| g11539.t1__seq_id=BLLR01000031.1_type=cds_species=Mincognita      | 239 | 96.3 | 46.1 | 24.7 | 29.7 | 26.4 |
| g4100.t1__seq_id=CEWN01000556.1_type=cds_species=Mjavanica        | 203 | 92.3 | 46.9 | 24.6 | 29.6 | 27.6 |
| g34890.t1__seq_id=CAJEWN010001880.1_type=cds_species=Menterolobii | 185 | 93.0 | 48.2 | 23.8 | 30.3 | 27.6 |
| g1731.t1__seq_id=JAEEAS010000058.1_type=cds_species=Marenaria     | 203 | 92.3 | 46.9 | 24.6 | 29.6 | 27.6 |
| g25165.t1__seq_id=CEWN01007035.1_type=cds_species=Mjavanica       | 182 | 91.3 | 45.7 | 24.7 | 30.2 | 26.9 |
| g3081.t1__seq_id=CAJEWN010000016.1_type=cds_species=Menterolobii  | 161 | 90.3 | 45.8 | 25.5 | 29.2 | 28.0 |
| g18725.t1__seq_id=BLLR01000069.1_type=cds_species=Mincognita      | 161 | 90.3 | 45.8 | 25.5 | 29.2 | 28.0 |
| g18693.t1__seq_id=BLLR01000069.1_type=cds_species=Mincognita      | 132 | 63.3 | 45.6 | 25.0 | 30.3 | 27.3 |
| g8399.t1__seq_id=CEWN01001434.1_type=cds_species=Mjavanica        | 116 | 61.0 | 45.4 | 25.0 | 31.0 | 25.0 |
+--------------------------------------------------------------+-------------+--------------------+------------------+---------+---------+---------+

Median of kmer hits to each target sequence: 203

dsRNA sense-arm sequence - 45.7% GC content
CGTTCAAATATGCTTGGGTGTTGGACAAGTTGAAGGCCGAGCGTGAACGTGGTATTACCATCGACATTGCTCTCTGGAAGTTCGAAACCGCCAAATACTATGTCACAATTATTGACGCTCCAGGACATCGTGACTTCATCAAGAACATGATTACTGGTACATCCC
AAGCTGATTGTGCTGTTTTGGTTGTTGCTTGTGGTACTGGAGAGTTCGAGGCTGGAATTTCTAAGAATGGTCAAACTCGCGAGCATGCTCTTCTCGCTCAGACTTTGGGAGTGAAGCAACTTATTGTTGCGTGTA
```

**Supplementary Figure 3: Universal *Meloidogyne* dsRNA with no off-target dataset.** 30 *Meloidogyne spp. TEF1a* transcript sequences were used as target inputs. All options were at the default settings (on-target *k*-mer length = 21, construct length = 300, 100 iterations). The final design had a median of 203 21 nt *k*-mer matchers to each target *TEF1a* sequence.

```
2024/12/10 13:55:50 dsRNAmax - dsRNA maximizer (Version: 1.1.12)
2024/12/10 13:55:50 Target FASTA File: ../ref/TEF.fa
2024/12/10 13:55:50 Off-target FASTA File: ../ref/GCF_000002985.6_WBcel235_genomic.fasta
2024/12/10 13:55:50 Loading target sequences...
     ---> 30 sequences loaded
2024/12/10 13:55:50 Applying bias modification to sequence 'g3046.t1__seq_id=CAJEWN010000016.1_type=cds_species=Menterolobii' at level 10...
2024/12/10 13:55:50 Getting target sequence kmers...
2024/12/10 13:55:50 8,902 target kmers loaded
2024/12/10 13:55:50 Removing off-target kmers from FASTA files...
Total off-target-matching kmers removed: 1305

2024/12/10 13:55:51 Finding best construct...

Results:
+----------------------------------------------------------------+-------------+--------------------+------------------+---------+---------+---------+
|                     TARGET SEQUENCE HEADER                     | 21NT MATCHES | SWG SIMILARITY (%) | KMER MEAN GC (%) | 5'U (%) | 5'A (%) | 5'C (%) |
+----------------------------------------------------------------+-------------+--------------------+------------------+---------+---------+---------+
| g3046.t1__seq_id=CAJEWN010000016.1_type=cds_species=Menterolobii | 236 | 97.3 | 44.8 | 25.8 | 28.4 | 27.1 |
| g17458.t1__seq_id=CAJEWN010000248.1_type=cds_species=Menterolobii | 167 | 94.0 | 42.7 | 29.3 | 25.1 | 30.5 |
| g4219.t1__seq_id=BLLR01000009.1_type=cds_species=Mincognita | 142 | 79.7 | 44.2 | 27.5 | 25.4 | 31.7 |
| g8398.t1__seq_id=CEWN01001434.1_type=cds_species=Mjavanica | 194 | 95.0 | 42.5 | 30.4 | 26.3 | 27.8 |
| g16315.t1__seq_id=CEWN01003635.1_type=cds_species=Mjavanica | 167 | 93.0 | 43.8 | 29.3 | 27.5 | 26.9 |
| g2430.t1__seq_id=JAEEAS010000070.1_type=cds_species=Marenaria | 194 | 95.0 | 42.5 | 30.4 | 26.3 | 27.8 |
| g7704.t1__seq_id=CEWN01001282.1_type=cds_species=Mjavanica | 176 | 89.3 | 45.2 | 27.8 | 25.0 | 27.8 |
| g26932.t1__seq_id=CEWN01007834.1_type=cds_species=Mjavanica | 194 | 95.0 | 42.5 | 30.4 | 26.3 | 27.8 |
| g2400.t1__seq_id=JAEEAS010000070.1_type=cds_species=Marenaria | 136 | 89.3 | 42.4 | 32.4 | 22.8 | 26.5 |
| g36955.t1__seq_id=CAJEWN010002398.1_type=cds_species=Menterolobii | 112 | 87.3 | 42.7 | 31.3 | 23.2 | 28.6 |
| g28756.t1__seq_id=CAJEWN010000898.1_type=cds_species=Menterolobii | 142 | 90.0 | 43.9 | 30.3 | 23.2 | 28.9 |
| g47475.t1__seq_id=JAEEAS010001409.1_type=cds_species=Marenaria | 194 | 95.0 | 42.5 | 30.4 | 26.3 | 27.8 |
| g11711.t1__seq_id=CEWN01002262.1_type=cds_species=Mjavanica | 189 | 94.0 | 46.4 | 25.9 | 25.9 | 26.5 |
| g9315.t1__seq_id=CAJEWN010000078.1_type=cds_species=Menterolobii | 218 | 96.0 | 46.1 | 25.7 | 27.5 | 25.7 |
| g17491.t1__seq_id=CAJEWN010000248.1_type=cds_species=Menterolobii | 131 | 87.7 | 44.1 | 29.8 | 24.4 | 26.7 |
| g1759.t1__seq_id=JAEEAS010000058.1_type=cds_species=Marenaria | 189 | 94.0 | 46.4 | 25.9 | 25.9 | 26.5 |
| g11578.t1__seq_id=BLLR01000031.1_type=cds_species=Mincognita | 218 | 96.0 | 46.1 | 25.7 | 27.5 | 25.7 |
| g4252.t1__seq_id=BLLR01000009.1_type=cds_species=Mincognita | 131 | 87.7 | 44.1 | 29.8 | 24.4 | 26.7 |
| g42043.t1__seq_id=CEWN01017646.1_type=cds_species=Mjavanica | 146 | 87.3 | 43.9 | 28.8 | 24.7 | 27.4 |
| g9360.t1__seq_id=CAJEWN010000078.1_type=cds_species=Menterolobii | 157 | 90.3 | 44.2 | 30.6 | 23.6 | 24.8 |
| g27345.t1__seq_id=JAEEAS010000752.1_type=cds_species=Marenaria | 146 | 87.3 | 43.9 | 28.8 | 24.7 | 27.4 |
| g11539.t1__seq_id=BLLR01000031.1_type=cds_species=Mincognita | 157 | 90.3 | 44.2 | 30.6 | 23.6 | 24.8 |
| g4100.t1__seq_id=CEWN01000556.1_type=cds_species=Mjavanica | 124 | 88.3 | 42.1 | 31.5 | 23.4 | 25.0 |
| g34890.t1__seq_id=CAJEWN010001880.1_type=cds_species=Menterolobii | 182 | 92.3 | 46.8 | 28.6 | 23.1 | 25.3 |
| g1731.t1__seq_id=JAEEAS010000058.1_type=cds_species=Marenaria | 124 | 88.3 | 42.1 | 31.5 | 23.4 | 25.0 |
| g25165.t1__seq_id=CEWN01007035.1_type=cds_species=Mjavanica | 124 | 88.3 | 42.1 | 31.5 | 23.4 | 25.0 |
| g3081.t1__seq_id=CAJEWN010000016.1_type=cds_species=Menterolobii | 124 | 88.3 | 42.1 | 31.5 | 23.4 | 25.0 |
| g18725.t1__seq_id=BLLR01000069.1_type=cds_species=Mincognita | 124 | 88.3 | 42.1 | 31.5 | 23.4 | 25.0 |
| g18693.t1__seq_id=BLLR01000069.1_type=cds_species=Mincognita | 243 | 97.3 | 44.4 | 26.3 | 28.8 | 25.9 |
| g8399.t1__seq_id=CEWN01001434.1_type=cds_species=Mjavanica | 194 | 95.0 | 42.5 | 30.4 | 26.3 | 27.8 |
+----------------------------------------------------------------+-------------+--------------------+------------------+---------+---------+---------+

Median of kmer hits to each target sequence: 162

dsRNA sense-arm sequence - 45.7% GC content
CGGAAGCTCGTTTTGAGGAAGTTAAGAATGAAGTCTCCAGCTTTATTAAGAAGATTGGTTACAACCCTGCAACTGTTGCGTTTGTCCCTATTTCTGGCTTTAATGGCGACAATATGTTGGAGCCGTCTGATAAGATGCCTTGGTTCAAAGGATGGGCTATTGAAA
GGAAGGATGGAAATGCTAGCGGGAAGACTTTGTTGGAAGCCCTTGACGCTATCCTGCCTCCAAGTAGGCCTACTGACAAGCCACTCCGACTTCCACTTCAAGATGTTTACAAGATTGGAGGTATTGGAACTGTGC
```

**Supplementary Figure 4: Biased *Meloidogyne* dsRNA with the *C. elegans* genome off-target dataset.** 30 *Meloidogyne spp. TEF1a* transcript sequences were used as target inputs. The *C. elegans* genome assembly (WBcel235) served as the off-target input in FASTA format, with the off-target cutoff set to 17. The bias header was g3046.t1, with a bias level of 10. All other options were at the default settings (on-target *k*-mer length = 21, construct length = 300, 100 iterations). The final design had a median of 162 21 nt *k*-mer matchers to each target *TEF1a* sequence.

**Supplementary Table 6: dsRNAmax performance benchmarks.** Desktop workstation with AMD Threadripper Pro 7985wx CPU, 512GB RAM. Where used, off-target length = 21nt

| Target organism/s | Number of input sequences | Size of input sequences | Off-target organism/s | Off-target data type | Size of off-target dataset | Time for dsRNA design |
|---|---|---|---|---|---|---|
| **4 root-knot nematode spp.** | 30 | 46,882nt | *C. elegans* | genome | 101MB | 5.9 seconds |
| **CMV virus isolates – RNA3** | 339 | 747,093 nt | None. | N/A | N/A | 5.9 seconds |
| **CMV virus isolates – RNA3** | 339 | 747,093 nt | Human, 4 honeybee species, monarch butterfly | transcriptome | 864MB | 6.5 seconds |
| **CMV virus isolates – RNA3** | 339 | 747,093nt | Cucumber rhizosphere metagenome sequencing (SRR31016968) | FASTQ files → .kmer using SeqToKmer (min kmer count = 2; 10min56seconds for conversion) | 7.7GB → 9.2GB 21nt kmers | 14.3 seconds |

**Supplementary Table 7: Comparison of dsRNAmax to currently active dsRNA design tools.** Non-accessible tools are not included[1].

| dsRNA design tool | Multiple target design | Off-target strategy | Off-target selection | Target optimisation strategy | Actively developed / maintained |
|---|---|---|---|---|---|
| **dsRNAmax** | Yes | Avoids any contiguous sense or antisense match of a set nucleotide length | Any sequence data in FASTA or .kmer format, from transcriptomes, genome, metagenome, read files and beyond. Amount limited only by computational resources | Maximising the number of dsRNA-derived kmers match individual target inputs | Yes |
| **dsCheck** | No | Checks for exact kmer matches, as well at 1nt and 2nt mismatches. Presents an 'off-target minimised' design | *D. melanogaster, C. elegans, A. thaliana*, *O. sativa* and *R. norvegicus* CDS only | Off-target minimisation focus | Yes – web server available |
| **siRNA-Finder** | No | Shows regions of a target transcript where off-target siRNAs are present | FASTA input creates Bowtie database. | Derived siRNA 5' nucleotide, siRNA end thermodynamic stability, target site accessibility | No longer maintained due to Python 2 legacy code. Binary available. Help documentation unavailable |

[1]Tools not accessible at the time of writing (2024) include e-RNAi (http://www.e-rnai.org forwards to https://e-rnai.dkfz.de/ - 404 page not found error), DEQOR (http://cluster-1.mpi-cbg.de/Deqor/deqor.html - Server Not Found), OfftargetFinder (http://rnai.specifly.org – the connection has timed out).
